# Diesel2p mesoscope with dual independent scan engines for flexible capture of dynamics in distributed neural circuitry

**DOI:** 10.1101/2020.09.20.305508

**Authors:** Che-Hang Yu, Jeffrey N. Stirman, Yiyi Yu, Riichiro Hira, Spencer L. Smith

## Abstract

Imaging the activity of neurons that are widely distributed across brain regions deep in scattering tissue at high speed remains challenging. Here, we introduce an open-source system with Dual Independent Enhanced Scan Engines for Large Field-of-view Two-Photon imaging (Diesel2p). Combining novel optical design, adaptive optics, and temporal multiplexing, the system offers subcellular resolution over a large field-of-view (∼ 25 mm^2^) with independent scan engines. We demonstrate the flexibility and various use cases of this system for calcium imaging of neurons in the living brain.

## Main

Two-photon microscopy^1^ has enabled subcellular resolution functional imaging of neural activity deep in the scattering tissue of mammalian brains^2^. However, conventional microscopes provide subcellular resolution over only small fields-of-view (**FOVs**), ∼ Ø0.5 mm. This limitation precludes measurements of neural activity distributed across cortical areas that are millimeters apart **(Fig. 1a)**. Obtaining subcellular resolution over a large FOV involves scaling up the dimensions of the objective lens and other optics, due to the Smith-Helmholtz invariant, also known as the optical invariant^3-5^. However, that is only half of the solution. Since high light intensities are required for efficient multiphoton excitation, two-photon imaging is typically implemented as a point-scanning approach, where an excitation laser beam is scanned over the tissue. Thus each voxel sampled entails a time cost, and the scan engine design constrains the time resolution^6^. Temporal multiplexing of simultaneously scanned beams can increase throughput^7^, and these can have either a fixed configuration^8-10^, or can be reconfigured during the experiment^11-14^. However, these simultaneously scanned, or “yoked”, multi-beam configurations strongly constrain sampling, because they preclude varying the scan parameters among the multiplexed beams. Optimal scan parameters (e.g., frame rate, scan region size) vary across distributed neural circuitry and experimental requirements, but yoked scanning requires using the same scan parameters for all beams. Therefore, a system featuring both a large imaging volume and independent multi-region imaging is needed, and can enable new experiments.

**Figure 1.**
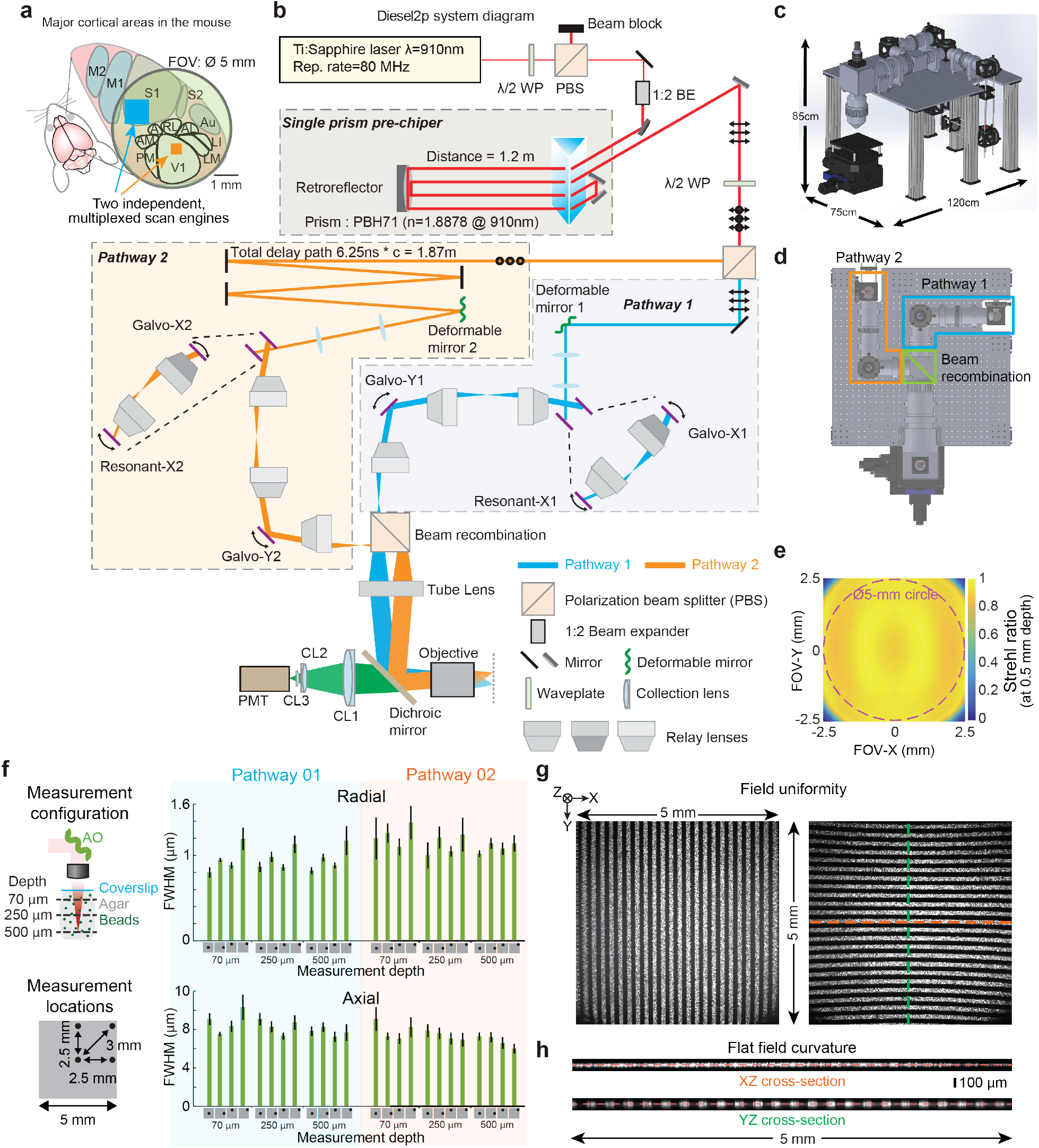
Diesel2p system features, layout, and performance benchmarks. (a) Functional cortical areas in the mouse brain are widely distributed. A field-of-view (FOV) of Ø5 mm can encompass multiple brain areas, and independent scan engines can to capture ongoing neural activity in multiple cortical areas simultaneously with optimized scan parameters. (b) Two imaging beams are temporally multiplexed and independently positioned in XY using two sets of resonant-galvo-galvo scan engines. First, overall power is attenuated using a half-wave plate (λ/2 WP) and a polarizing beam splitting cube (PBS). A 2X beam expander (1:2 BE) enlarges the beam for the clear aperture of the deformable mirrors adaptive optics (AO). A custom single-prism pre-chirper offsets system dispersion to maintain transform-limited pulses at the focal plane. A second λ/2 WP and PBS pair divides the beam into two pathways. Pathway 2 (s-polarization) passes to a delay arm where it travels 1.87 m further than Pathway 1 using mirrors, thus delaying it by 6.25 ns relative to Pathway 1 (p-polarization). Both pathways each proceed to deformable mirrors for adjusting the focal plane and correcting optical aberrations before being directed to resonant-galvo-galvo scan engines. All scanning mirrors are optically relayed to each other. Each pathway then passes through a scan lens before being combined with a beam recombination relay. A tube lens and an infrared-reflective dichroic mirror relay the two multiplexed beams onto the back aperture of the objective. Fluorescence (green) is directed to a photomultiplier tube (PMT) via an assembly of collection lenses (CL1, CL2, CL3). (c) An oblique view of a 3-D model of the system and its footprint. (d) A top view with the arrangement of the two scan engines highlighted. (e) Plot of the model Strehl ratio across the scan area indicates diffraction limited performance (Strehl ratio > 0.8) across ∼ 25 mm^2^. (f) Multiphoton excitation PSF measurements were made with subresolution beads in agar under a coverslip at three depths and four locations, for both of the AO-equipped, temporally multiplexed beam pathways. Full width at half-maximum (FWHM) of the Gaussian fits (mean ± S.D.) are plotted. (g) XY images of a calibrated, structured fluorescent sample with a periodic line pattern (5 lines per millimeter) in two orientations acquired under the full scan range of the system. Each image shows 25 lines on the top edge (left image) and on the left edge (right image), receptively, verifying a 5 × 5mm FOV. (h) The XZ image along the orange dashed line and the YZ image along green dashed line in (g) are also plotted. The imaging pattern is colinear with the straight lines, suggesting a flat field both in x and y directions across the FOV.

To address this need, we developed a system with a large FOV, subcellular resolution, and dual independent scan engines for highly flexible, asymmetric multi-point sampling of distributed neural circuitry. Here, we present a custom two-photon system with dual scan engines that can operate completely independently. Each arm has optical access to the same large imaging volume (> Ø5 mm FOV) over which subcellular resolution is maintained in scattering tissue to typical 2-photon imaging depths. These two arms use adaptive optics (**AO**) for wavefront shaping, temporal multiplexing for simultaneous imaging, and polarization optics for beam recombination. Due to the independence of the arms, the input lasers can come from the same source or different sources (multi-wavelength). Moreover, each arm can use multiple sources simultaneously, for example, in imaging and photoactivation experiments. We refer to the system as the **Diesel2p** (Dual Independent Enhanced Scan Engines, Large-Field Two-Photon).

The Diesel2p has two major design features. First, it has a > Ø5 mm FOV to encompass multiple cortical areas and provides subcellular resolution throughout **(Fig. 1a)**. Second, the Diesel2p can perform simultaneous two-region imaging using two scan engine arms. In contrast to prior work^11,13^, these two arms are completely independent. They can each scan any region and be configured to with different imaging parameters (e.g., pixel dwell time, scan size) including random access scanning **(Fig. 1b-d)**. To achieve these two features, several scan engine components were custom designed and manufactured: the optical relays, the scan lens, the tube lens, and the objective. The full optical prescriptions are provided in this report **(Supplementary Fig. 1-5**, ZEMAX models). The system was optimized as a whole, rather than optimizing components individually, to minimize the aberrations across scan angles up to ± 5 degrees at the objective back aperture and primarily for the excitation windows of 910 ± 10 nm and 1050nm ± 10 nm. The optics use an infinity-corrected objective design to facilitate modifications and modularity. Based on the optical design model, the Strehl ratio exceeded 0.8 (consistent with a diffraction-limited design) over an area only slightly smaller than a 5 × 5 mm^2^ (25 mm^2^) square **(Fig. 1e)**.

The Diesel2p system uses two independent scan engines to access two areas simultaneously **(Fig. 1a**,**b**,**d)**, as opposed to one beam jumping back and forth between two areas sequentially. In the sequential imaging regime, information is missed both during the scanning of the other area and during the time of jumping. This latter dead time is a larger fraction of the duty cycle as the frame rate increases **(Supplementary Fig. 6)**. The laser beam, after passing through the dispersion compensator^15^, is split into two pathways by a polarization beam splitter **(Fig. 1b)**. The temporal multiplexing is set by delaying one laser beam’s pulses relative to the other (for an 80 MHz system as used here, the delay is 6.25 ns). Beams are guided into two independent scan engines. Each scan engine consists of an x-resonant mirror, an x-galvo mirror, and a y-galvo mirror in series, each at conjugate planes connected by custom afocal relays. The x-resonant mirror provides rapid and length-variable x-line scanning, up to 1.5 mm. The x-and y-galvo mirrors provide linear transverse scanning across the full FOV. Therefore, each arm of the scan engine can arbitrarily position the imaging location within the full FOV, and scan with parameters that are completely independently from the other arm. Each pathway is also equipped with a deformable mirror AO for both rapidly adjusting the focal plane axially **(Supplementary Fig. 7a, Supplementary Video 1)** and correcting optical aberrations **(Supplementary Fig. 7b, c and Supplementary Video 2)**.

Next, we measured the resolution of the Diesel2p system by taking z-stacks of 0.2-µm fluorescent beads at various positions and depths. The lateral and axial resolutions were estimated from the full-width-at-half-maximum (FWHM) of measured intensity profiles. For beads at each XYZ location in the FOV, measurements were made with the deformable mirror optimized to act as an AO element. Overall, the lateral FWHM was ∼1 µm and axial FWHM was ∼8 µm across the 5-mm FOV and up to 500-µm imaging depth **(Fig. 1f)**. This indicates a space-bandwidth product of ∼ (25 mm^2^ / 1 µm^2^) = 25 * 10^6^, or 25 megapixels. The use of the deformable mirror as an AO element reduced the variation across the measured volume, and improved the axial resolution by ∼2 µm **(Supplementary Fig. 8)**. The AO also enabled imaging of neural activity over 3.5 mm from the center of the FOV, implying a 7-mm diameter along the diagonal axis (**Supplementary Fig. 7b**). These results show that the Diesel2p system maintains subcellular resolution over an imaging field larger than a Ø5 mm circle, allowing measurement of neural activity over at least a 25 mm^2^ area.

For efficient multiphoton excitation, the characteristics of the ultrafast pulses must be maintained over the full FOV. We measured pulse characteristics at the focal plane using the frequency-resolved optical gating (**FROG**) technique^16^. Throughout the FOV, the pulse width is maintained at ∼110 fs, and the pulse front tilt and the spatial chirp remain low (**Supplementary Fig. 9**). Together with the bead measurement, these results show that the Diesel2p system has consistent resolution and spatiotemporal pulse characteristics across the entire FOV.

We next verified the imaging FOV by imaging a structured fluorescent sample with repetitive 5 lines per mm (57-905, Edmund Optics). Images contain 25 lines along both the x and y directions, indicating a 5-mm length on each axis of the FOV **(Fig. 1g)**. The result demonstrates that the Diesel2p system has a FOV larger than Ø5-mm circle and close to a 5 × 5 mm^2^ field, consistent with the nominal model performance **(Fig. 1e)**. By z-scanning through this sample and rendering the x-z and y-z cross-section, we found that the thin fluorescence pattern is nearly co-linear with a straight line, indicative of a field curvature < 30 µm over 5000 µm of FOV **(Fig. 1h)**. This result indicates that the Diesel2p system has a flat field, minimizing the difference in depths between the center and the periphery of the FOV.

After benchmarking the optical performance imaging system, we performed a series of *in vivo* imaging experiments with neurons expressing the genetically encoded calcium indicator, GCaMP6s^17^. The brain tissue under a 5-mm diameter cranial window was positioned within the 5 × 5 mm^2^ FOV, and subcellular detail was resolved in individual neurons across the FOV **(Fig. 2a)**. This result confirms that the Diesel2p system preserves subcellular resolution throughout the FOV. Next, we positioned the two pathways to image two adjacent stripes of cortex simultaneously and set each pathway to scan a large FOV of 1.5 mm x 5 mm (1024 × 4096 pixels) **(Fig. 2b and Supplementary Video 3)**. Calcium signals from 5,874 neurons were detected from these two stripes **(Fig. 2c)**. The raw calcium signals had a signal-to-noise ratio of 7.9 ± 2.5 (mean ± standard deviation) **(Fig. 2d)**, and this supported robust spike inference **(Fig. 2e)**, which we used to calculate the correlation matrix **(Fig. 2f)**. In this data set, we imaged a total area of 15 mm^2^ with a pixel resolution of ∼ 1.5 × 1.2 µm^2^ (undersampling the resolution), and an imaging rate of 3.84 frames/s, resulting in a pixel throughput of 32.3 megapixels/s over 15 mm^2^. This data set demonstrates the ability of the system to measure neuronal activity with high fidelity over a large FOV.

**Figure 2.**
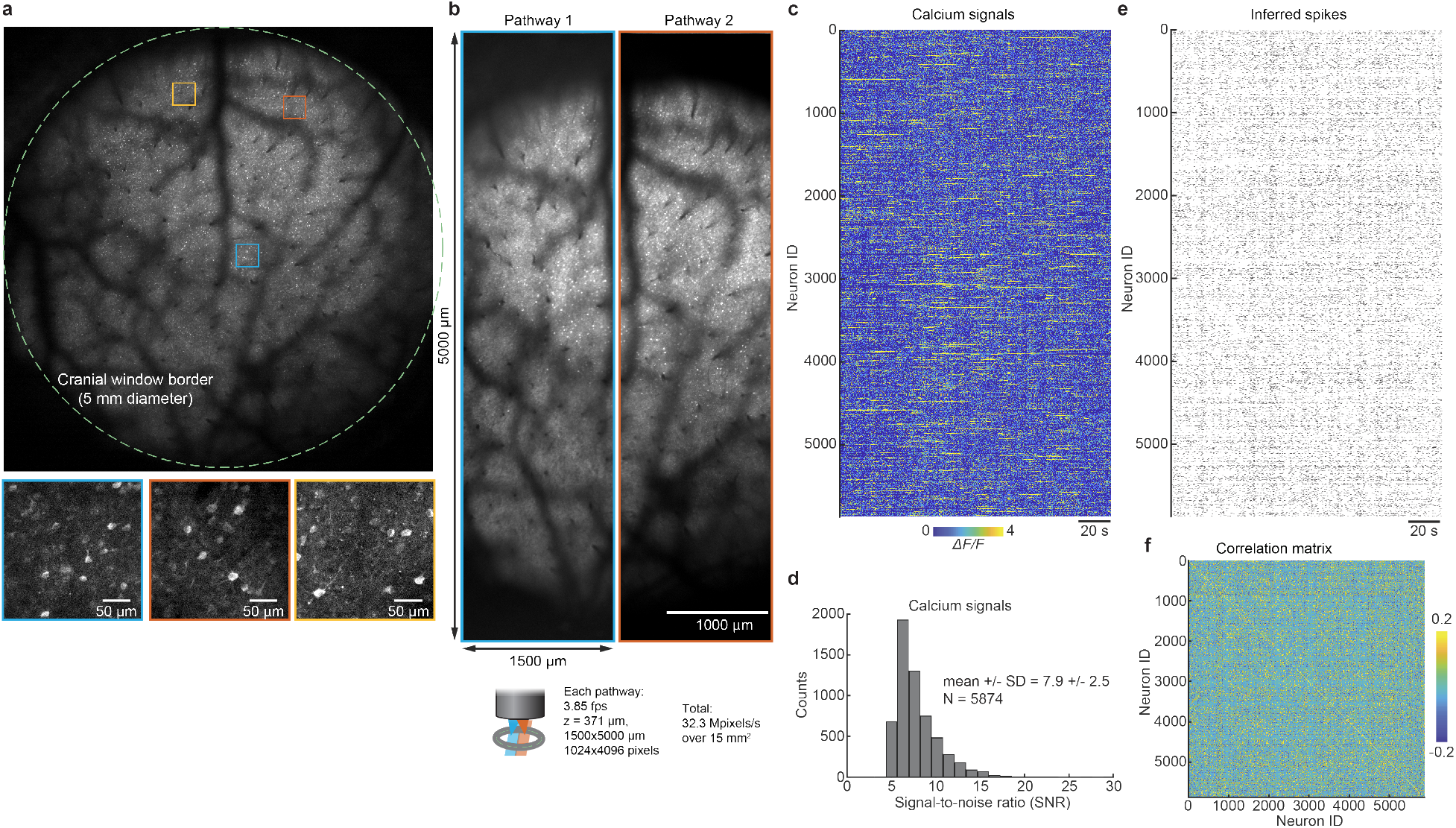
Diesel2p provides subcellular resolution two-photon imaging of neural activity across its FOV. (a) Diesel2p’s galvo-galvo raster scanning records neuronal activity at the depth of 345 µm through a cranial window with a diameter of 5 mm (dashed line) on a transgenic mouse expressing the genetically encoded fluorescent calcium indicator GCaMP6s in excitatory neurons. Three zoom-in views from different sub-regions (colored squares) show the preservation of the subcellular resolution. (b) Neuronal activity in two strips of cortex are recorded using Diesel2p’s two pathways simultaneously (blue and orange rectangles), covering a combined area of 3 mm x 5 mm with a frame rate of 3.85 frames/s. The imaging depth is 371 µm. The pixel number for each image is 1024 × 4096 pixels. (c) In this data set, calcium signals were recorded from 5,874 active neurons. (d) The histogram of the transients’ signal-to-noise ratio in (c). (e) Ca^2+^ signals in (c) were used to infer spike times. (f) Spikes in (e) were used to measure > 17 million cross-correlations the population.

To demonstrate the flexibility of the Diesel2p system, we performed four test experiments. First, we imaged two regions that were 4.36 mm apart, which is equivalent to the distance between the primary visual cortex and the motor cortex. Moreover, we configured the imaging fields, frame acquisition speeds, and the pixel numbers independently for the two regions **(Fig. 3a and Supplementary Video 4)**. Non-multiplexed imaging with these parameters would reduce the acquisition rate and involve > 20% dead time **(Supplementary Fig. 6)**, and yoked multiplexing would require a compromise in imaging parameters so that both regions would have had the same size and scan rate. Thus, the Diesel2p system enables new classes of measurements. Second, to demonstrate that the two pathways can be overlapped, we set Pathway 1 to image a subregion of the region imaged by Pathway 2 **(Fig. 3b and Supplementary Video 5)**. Third, we increased the number of imaging regions within a single imaging session by repositioning each of the pathways between two sub-regions. To scale up the number of regions, each of the pathways alternated (every other frame) between two sub-areas at different depths. A total of four sub-areas that differed in XY locations and Z depths are imaged at the same time **(Fig. 3c and Supplementary Video 6)**. Fourth, we used random-access scanning of cell bodies in conjunction with large FOV imaging. We configured Pathway 1 to raster scan a 2.25 mm^2^ area, and Pathway 2 executed a random-access scan of 12 cell bodies **(Fig. 3d and Supplementary Video 7)**. Together, these results, enabled by both the large FOV and the independent multiplexed scanning, demonstrate the flexibility of the Diesel2p system to enable new measurements and experiments.

**Figure 3.**
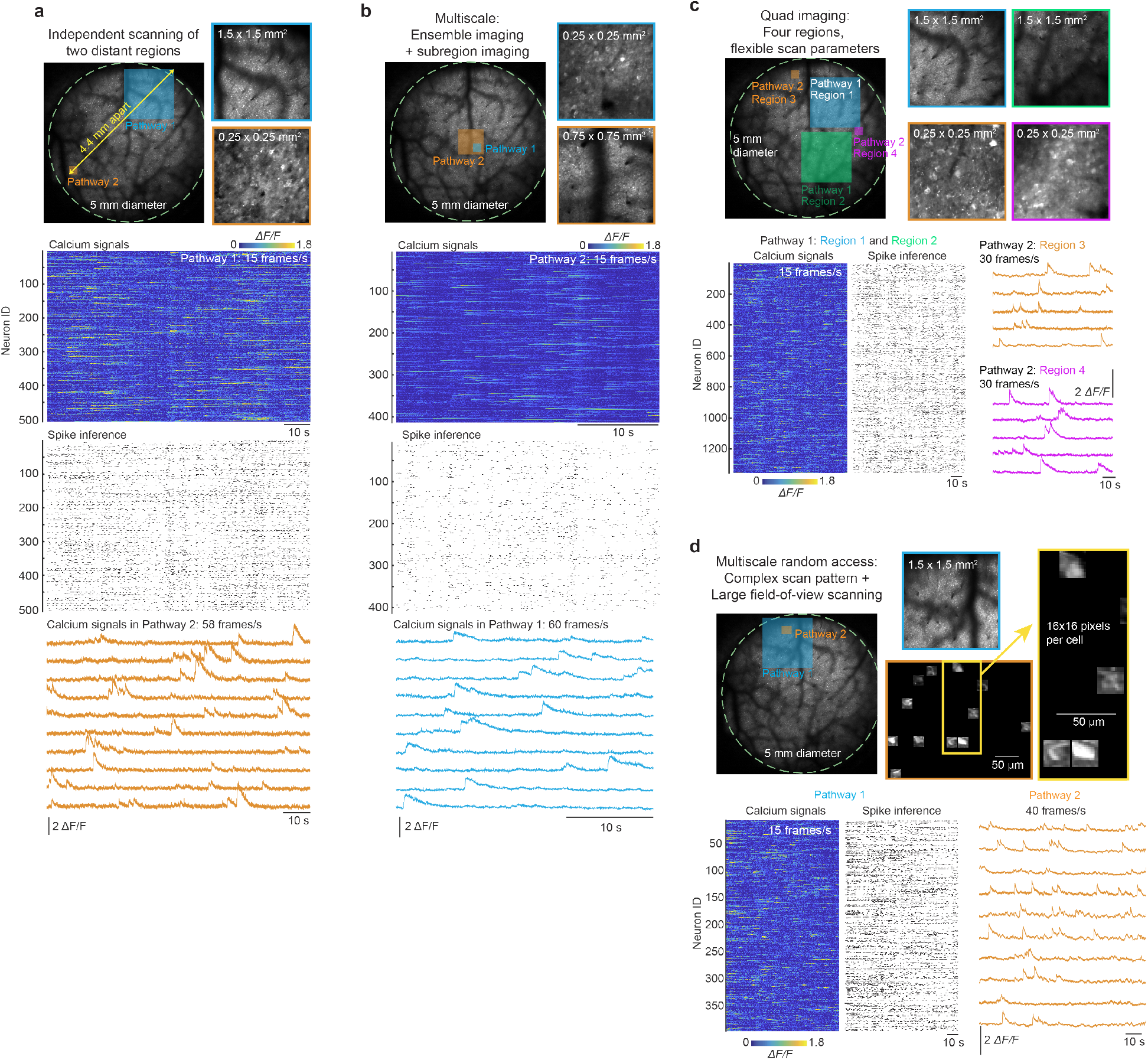
Flexible measurement with dual independent scan engines. (a) Neuronal activity at two distant regions (4.36-mm apart) are imaged simultaneously with independent imaging parameters. The blue and orange boxes indicate the imaging sizes and positions within the full FOV. Expanded regions are shown at right. Calcium signals from 500 neurons imaged in Pathway 1 were used to infer spikes. Simultaneously, Pathway 2 imaged calcium signals at 58 frames/s 4.36 mm away. (b) Neuronal activity from two overlapped regions are imaged simultaneously with different imaging parameters. Over 400 neurons were imaged in Pathway 2 while neural activity was imaged at 60 frames/s in Pathway 1. (c) Neuronal activity from four separate regions are imaged with two pathways independently positioned and then repositioned within the full FOV. By serially offsetting the galvo scanners, Pathway 1 accesses the blue and green regions, and Pathway 2 accesses the orange and the magenta regions. By changing the curvature of the AO, Pathways 1 and 2 also image at different depths. Calcium signals and inferred spike trains from neurons in the Pathway 1 data (Region 1 and Region 2 combined) are shown. Example calcium signals are shown for neurons from Region 3 (orange) and Region 4 (magenta) scanned by Pathway 2. (d) A combination of raster scanning and random-access scanning is configured on Pathways 1 and 2 for neuronal imaging. While Pathway 1 performs raster scanning, Pathway 2 performs random-access scanning for 12 cell bodies. Calcium signals and the inferred spike trains are shown for neurons imaged by Pathway 1. Example calcium signal traces are shown for neurons imaged by Pathway 2 (orange).

In summary, we present a newly developed imaging system to enable flexible, simultaneous, multi-region, multiphoton excitation in scattering tissue. The Diesel2p system has subcellular resolution over a ∼ 25 mm^2^ FOV with very low field curvature, linear galvo access to the full FOV, and resonant scan size of 1.5 mm. These optics and scan specifications are combined with a novel layout of two independent scan engines, each with deformable mirrors for fast z-focus and aberration correction. The temporally multiplexed imaging pathways can record neural activity in two arbitrarily selected portions of the imaging volume simultaneously.

This new system is also designed to facilitate experiments with behaving animals. The objective lens is air immersion, so no water interface is required, and it has an 8-mm-long working distance, to enable a variety of headplate designs and other instrumentation that needs to be close to the imaged area (e.g., electrode arrays). In addition, the objective can rotate 360 degrees. The advantage of an air immersion objective is evident when the objective is fixed at an angle for imaging the brain of a behaving animal from the side. These ergonomic features can facilitate behavior experiments that require flexibility for animal posture, further increasing the flexibility of the Diesel2p system.

The Diesel2p system uses an infinity-corrected objective design, making it compatible with a range of extensions to multiphoton imaging, including Bessel-beam scanning^18^ and reverberation microscopy^19^ to enhance the volumetric imaging capability. It can also be combined with two-photon optogenetics approaches to perform simultaneous multiphoton neuronal imaging and functional perturbation^20^. Wavelength multiplexing can be implemented to add another pulsed laser (e.g. a 1040 nm pulsed laser) into the Diesel2p system, enabling the dual-color imaging of two different molecules simultaneously **(Supplementary Fig. 10)**. The Diesel2p system enables new measurements of neural activity, is compatible with a range of variants of multiphoton imaging, and is a fully documented and open optical design that can be extended to support studies of neural interactions across brain areas.

---

**Supplementary Figure 1.**
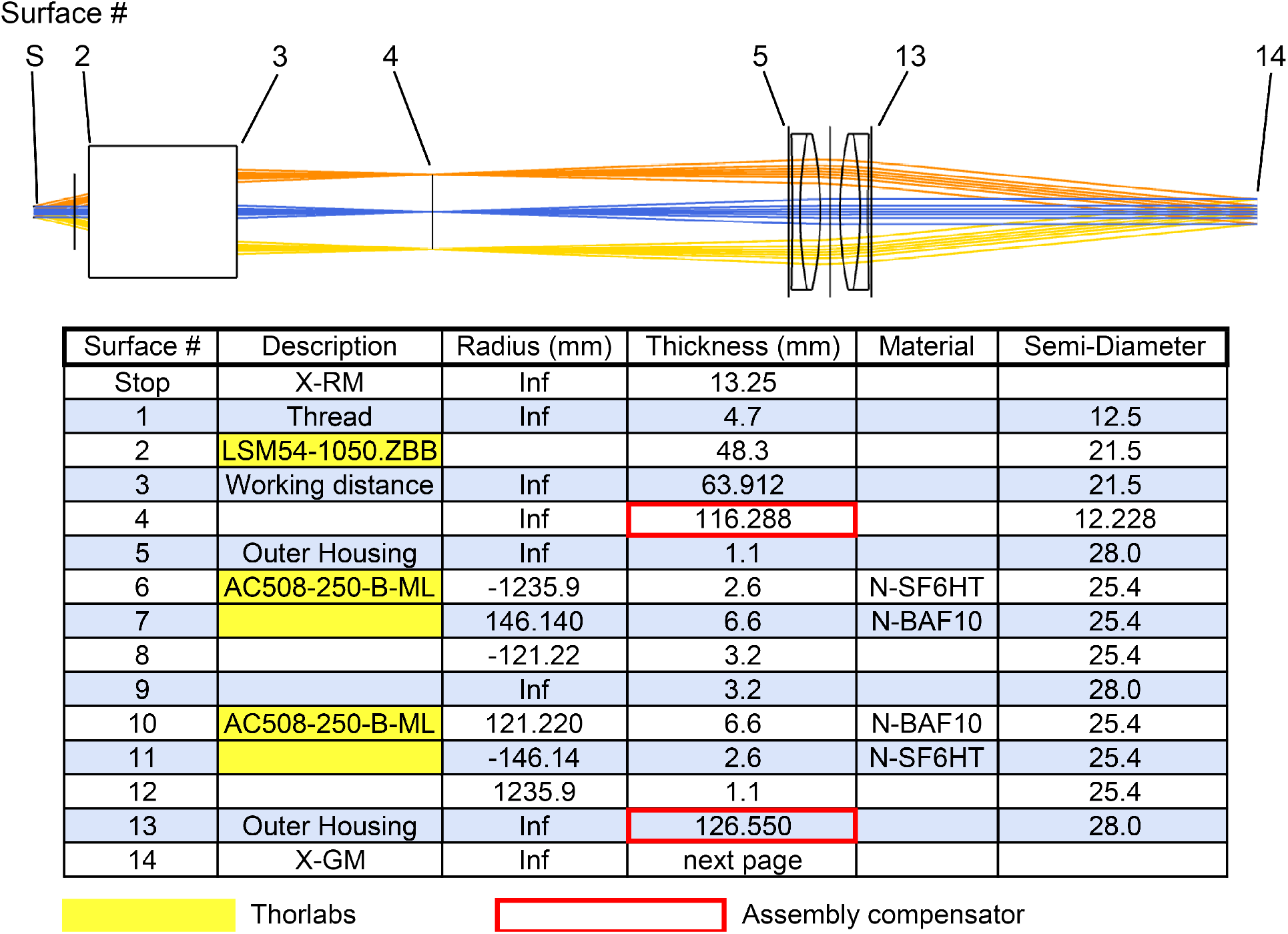
Full prescription data for X-resonant mirror (X-RM) to X-galvo mirror (X-GM) relay. This optical relay was constructed using commercial off-the-shelf (COTS) components from Thorlabs. The red outlined axial separation are used as compensators in system assembly.

**Supplementary Figure 2.**
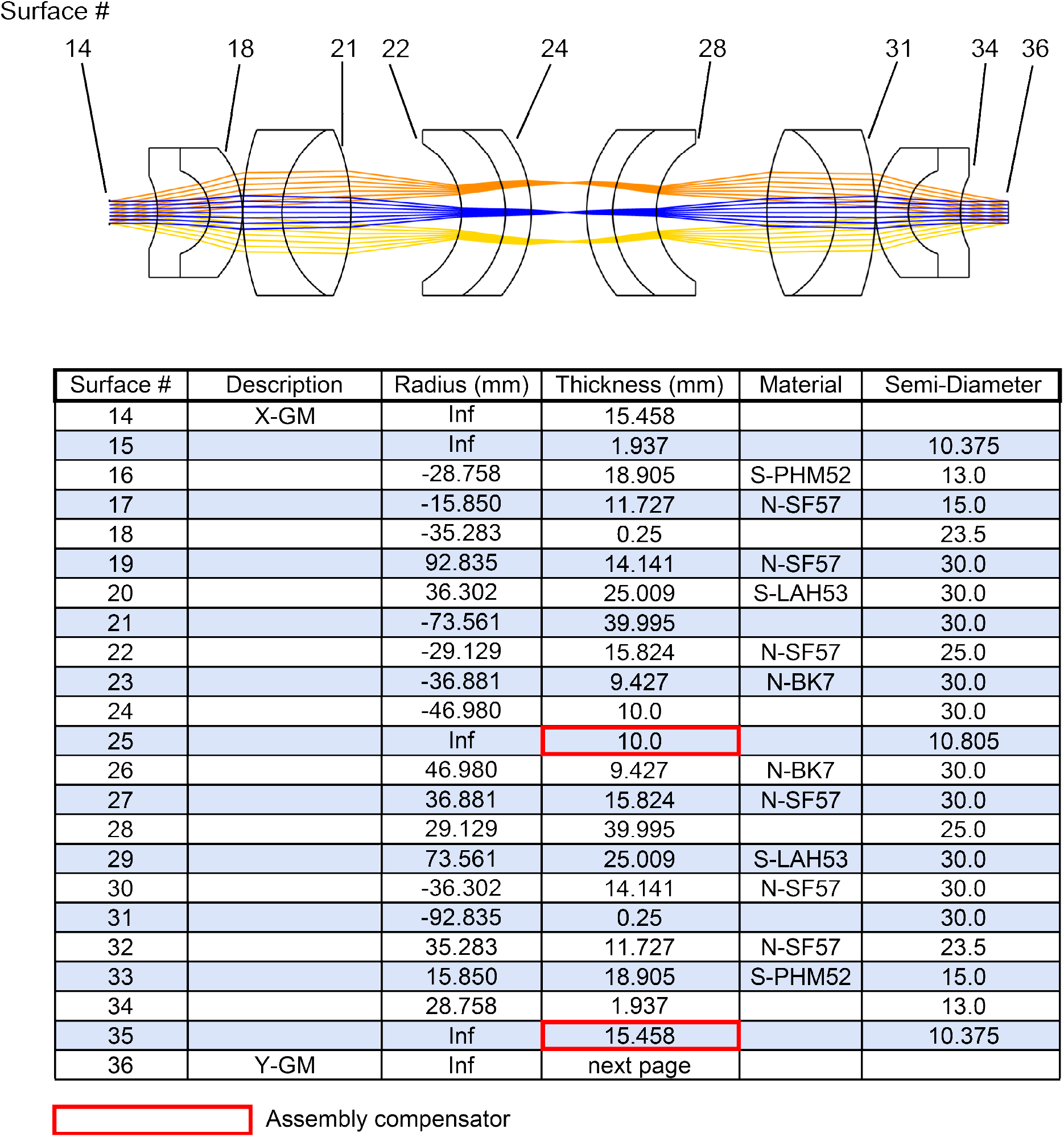
Full prescription data for X-galvo mirror (X-GM) to Y-galvo mirror (Y-GM) relay. The red outlined axial separation are used as compensators in system assembly.

**Supplementary Figure 3.**
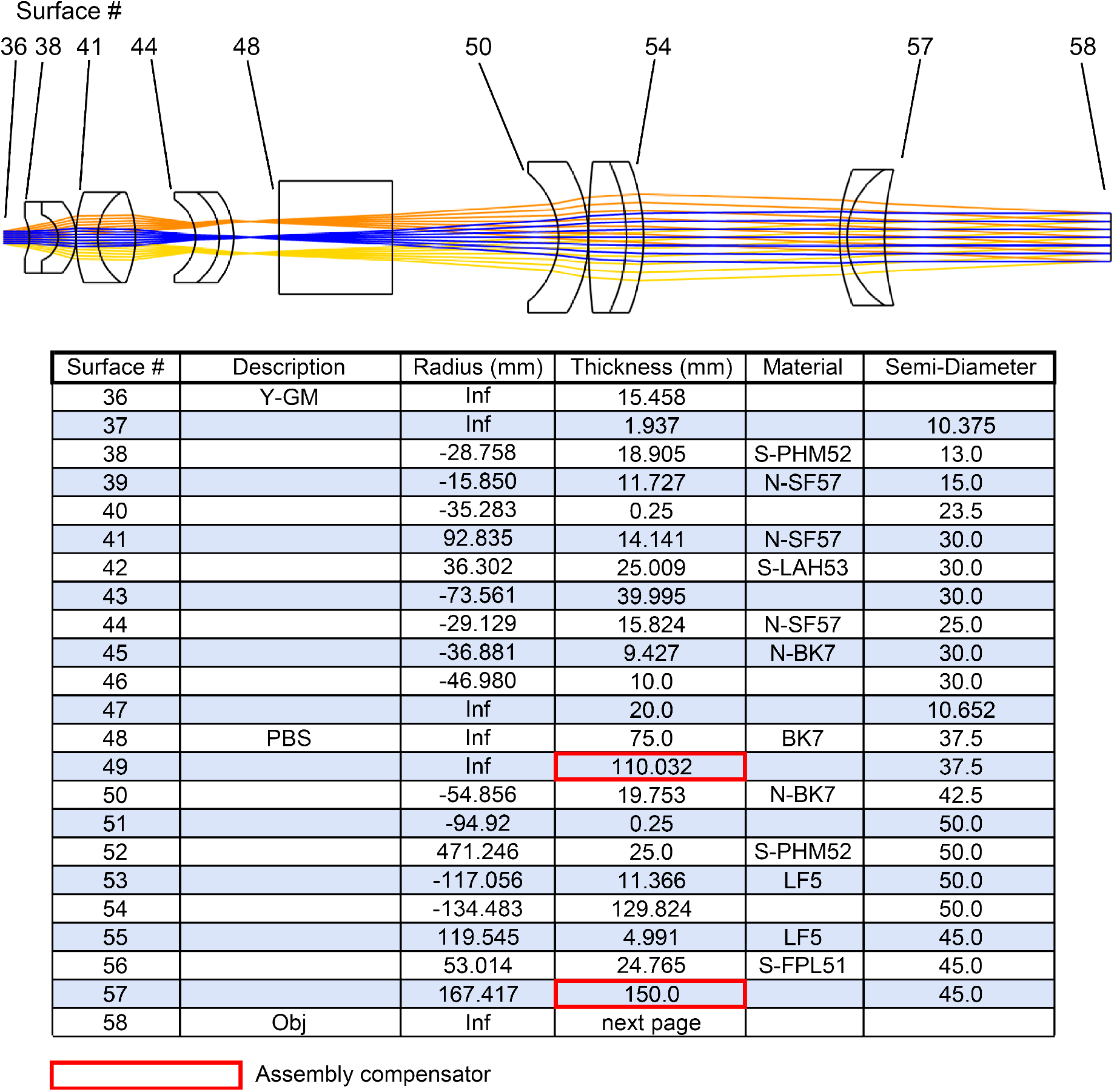
Full prescription data for system from the Y-galvo mirror (Y-GM) to the objective. The polarizing beam splitter (PBS) was offset from the focal point of the relay to minimize photo-damage to the optical cement. The terminal lens in this system has a 90 mm diameter in order to minimize vignetting at the extreme scan angles. The red outlined axial separation are used as compensators in system assembly.

**Supplementary Figure 4.**
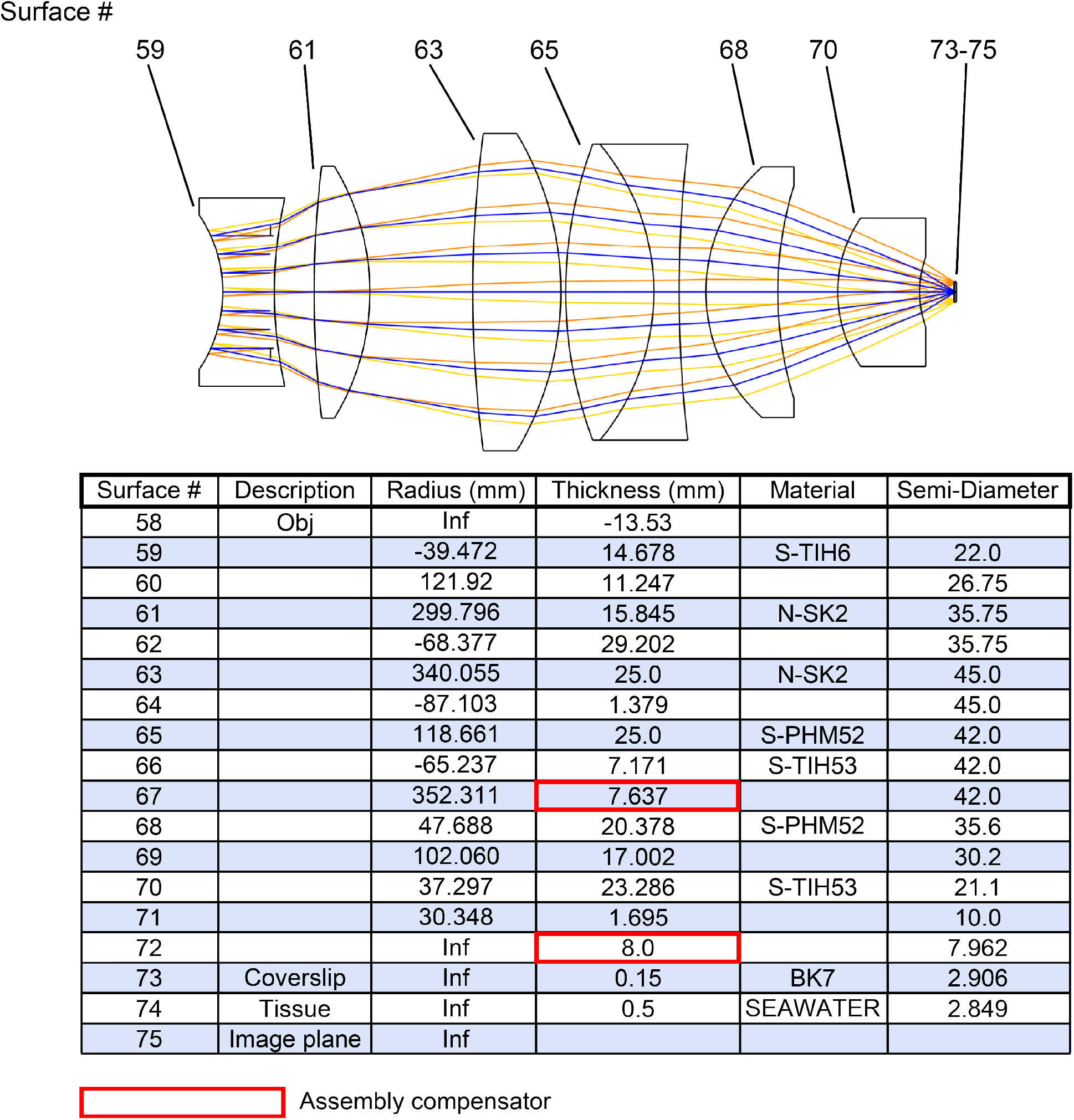
Full prescription data for an infinity-corrected, f = 30 mm, NA = 0.54 objective. The objective has ∼8 mm of working distance and 30-mm focal length. The axial separation on surface 67 (red outline) is used as a compensator in the objective assembly. Focus compensation occurs at surface 72.

**Supplementary Figure 5.**
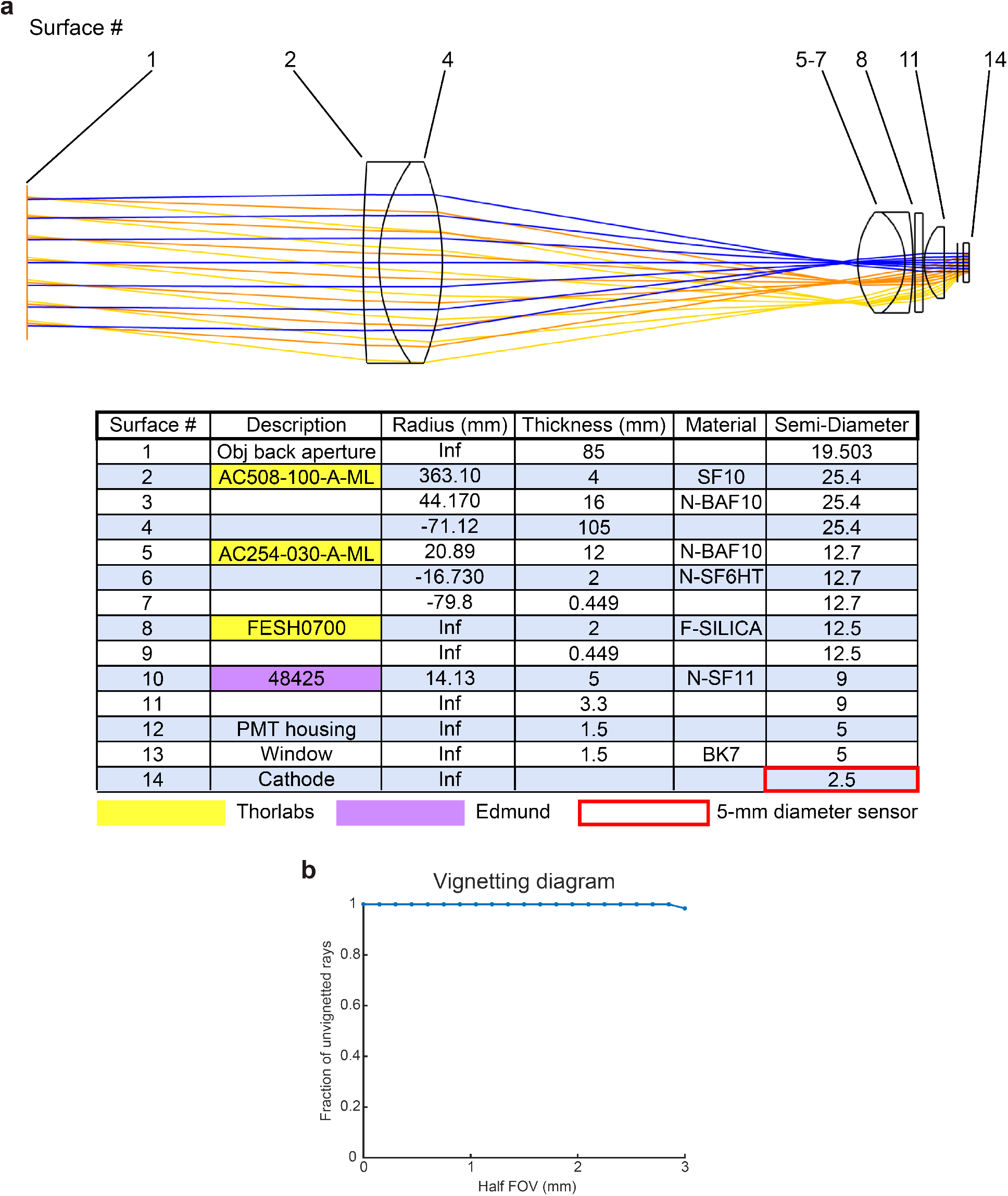
Full prescription data and vignetting diagram for the collection relay. (a) This optical relay was constructed using COTS components from Thorlabs and Edmund. (b) The collection efficiency is nearly 100% up to a 3-mm half FOV (6-mm full FOV) with a sensor diameter of 5 mm.

**Supplementary Figure 6.**
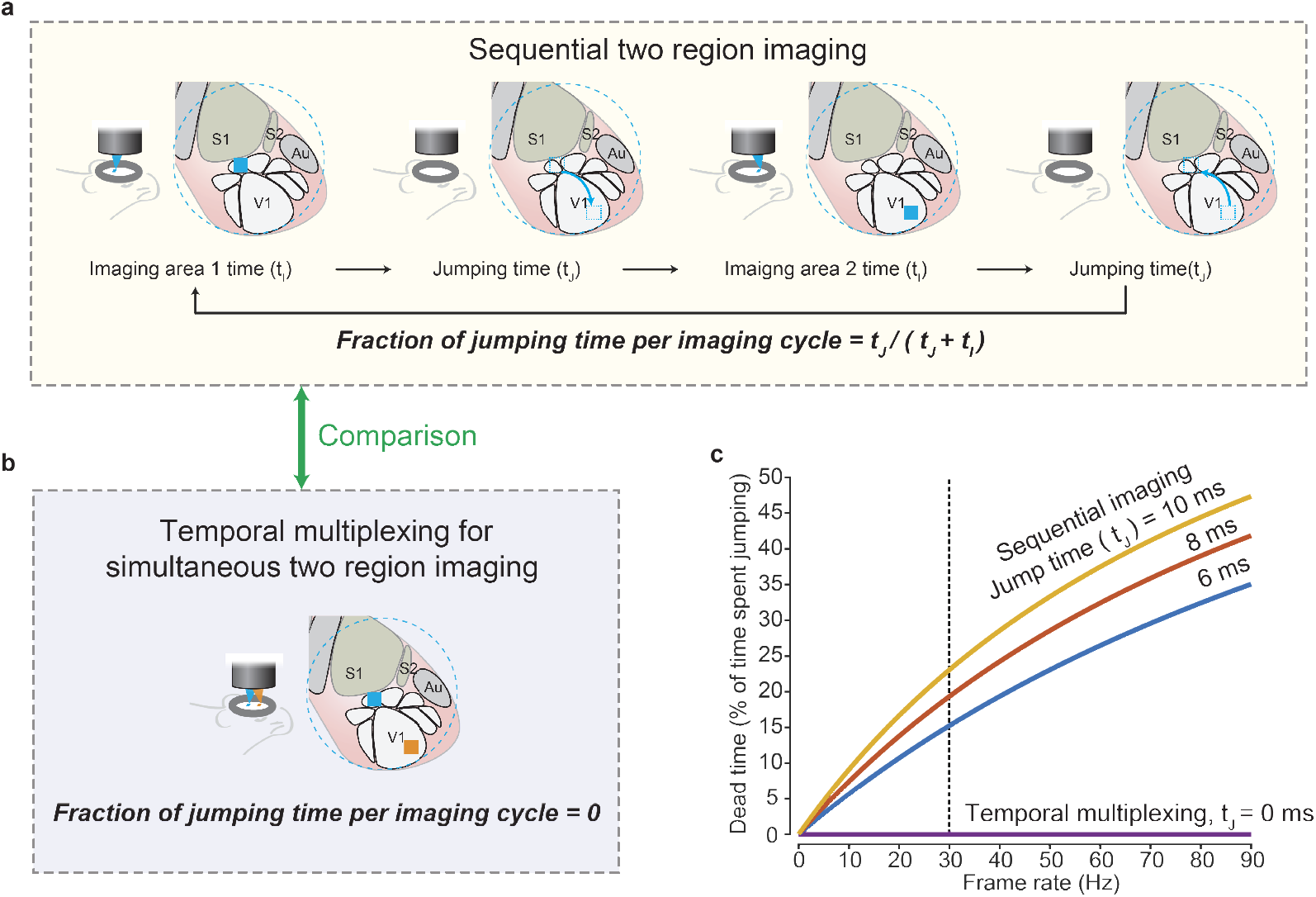
Zero jumping time for simultaneous two-region imaging. (a) The cartoon shows the pipeline of the sequential imaging approach. Between two imaging acquisitions, a period of time is taken for the scanners to jump from a position to the next position. During an imaging cycle, a fraction of time is spent on the jumping. (b) Zero fraction of time is spent on the jumping for temporal multiplexing imaging approach. (c) A chart shows the fraction of jumping time versus the frame rate for the sequential imaging and the temporal multiplexed imaging. The higher the frame rate is, the larger fraction of the jumping time is for sequential imaging. The vertical dashed line indicated the 30-Hz frame rate, frequently referred to as real-time frame rate.

**Supplementary Figure 7.**
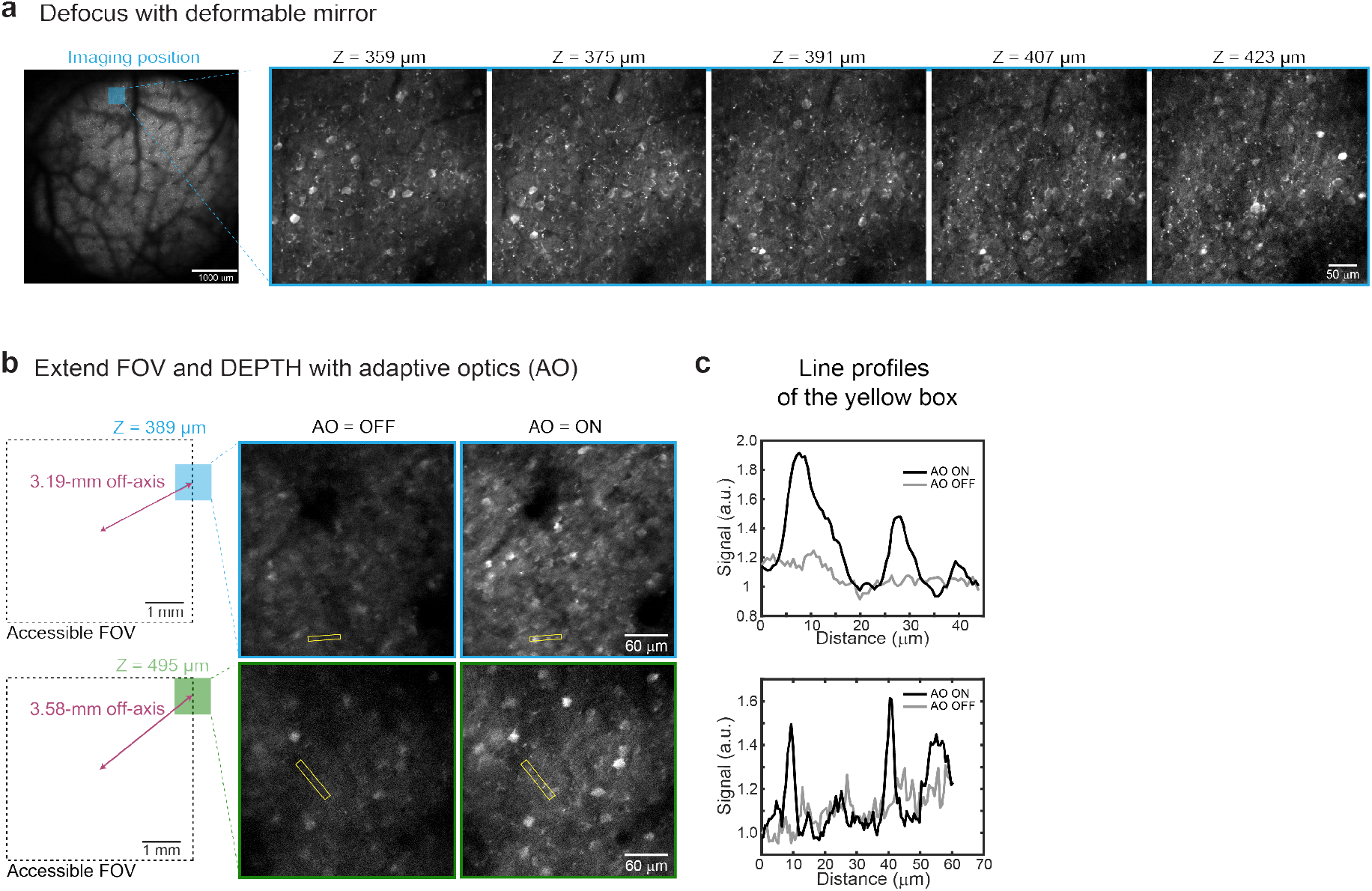
Deformable mirrors extend the functionality and FOV. (a) Neuronal activity imaged at different depths by changing AO curvature. The lateral imaging position (blue) is shown on the full FOV (left). Images at each z plane is shown (right). (b) Images measured at off-center positions >3 mm in the depths of 389 µm (blue) and 495 µm (green) without and with the AO correction. The image contrasts are set the same. (c) Signal profiles in the yellow boxes in (b). Profiles are normalized to the noise level.

**Supplementary Figure 8.**
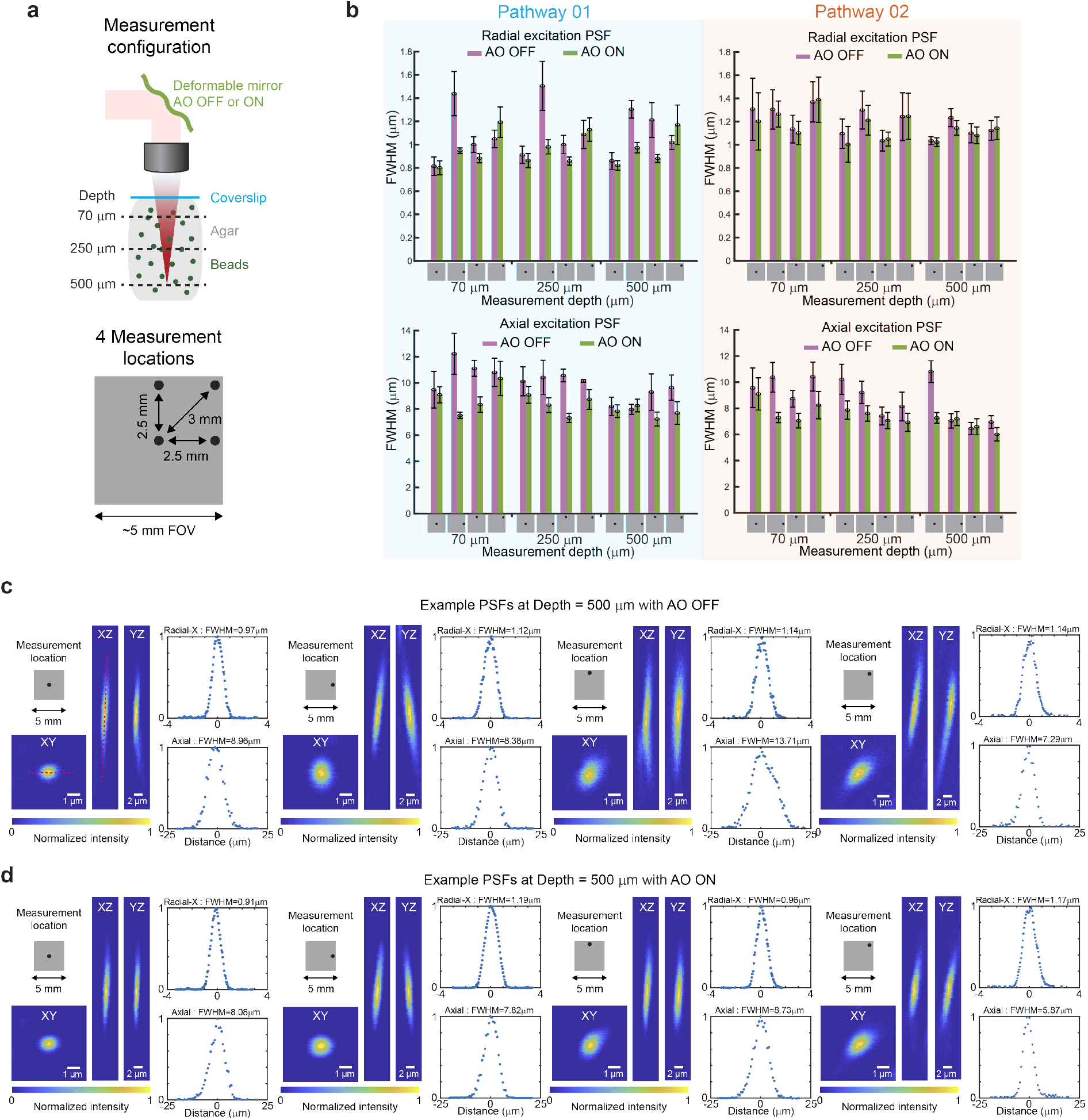
Complete point spread function characterization of Diesel2p. (a) 0.2 µm fluorescent beads were embedded in 0.75% agarose gel. 40 µm z-stacks were acquired, each centered at one of three depths (70 µm, 250 µm, 500 µm). At each depth, beads at four lateral locations were measured: on axis, 2.5-mm off-axis near the two edges, and 3-mm off-axis diagonally near the corner of the FOV. At each position, measurements were done with the AO flattened (OFF) or deformed (ON). (b) A complete summary of the excitation PSF measurements at positions indicated in (a) for both of the temporally multiplexed beam pathways with AO OFF and ON. FWHM of the Gaussian fits ± the s.d. for measurements from eight different beads radially and axially are calculated. The radial PSF is the average of both the radial FWHMs in the X and Y directions. (c) Radial and axial excitation PSF volume measurements were made at the indicated locations and the depth of 500 µm where the example images are shown from the XY, XZ, and YZ cross-sections, respectively. The intensity profiles of the beads (red lines) in the X direction on the XY plane and in the Z direction are plotted, which are fitted to a Gaussian curve to extract the radial-X and axial full-width-half-maximum (FWHM) of the PSF. This row shows the measurements as the AO was set flat. (d) This row shows the same z-stack measurement from exact the same bead in (c) after the AO was applied and configured to maximize the fluorescence signal from that bead.

**Supplementary Figure 9.**
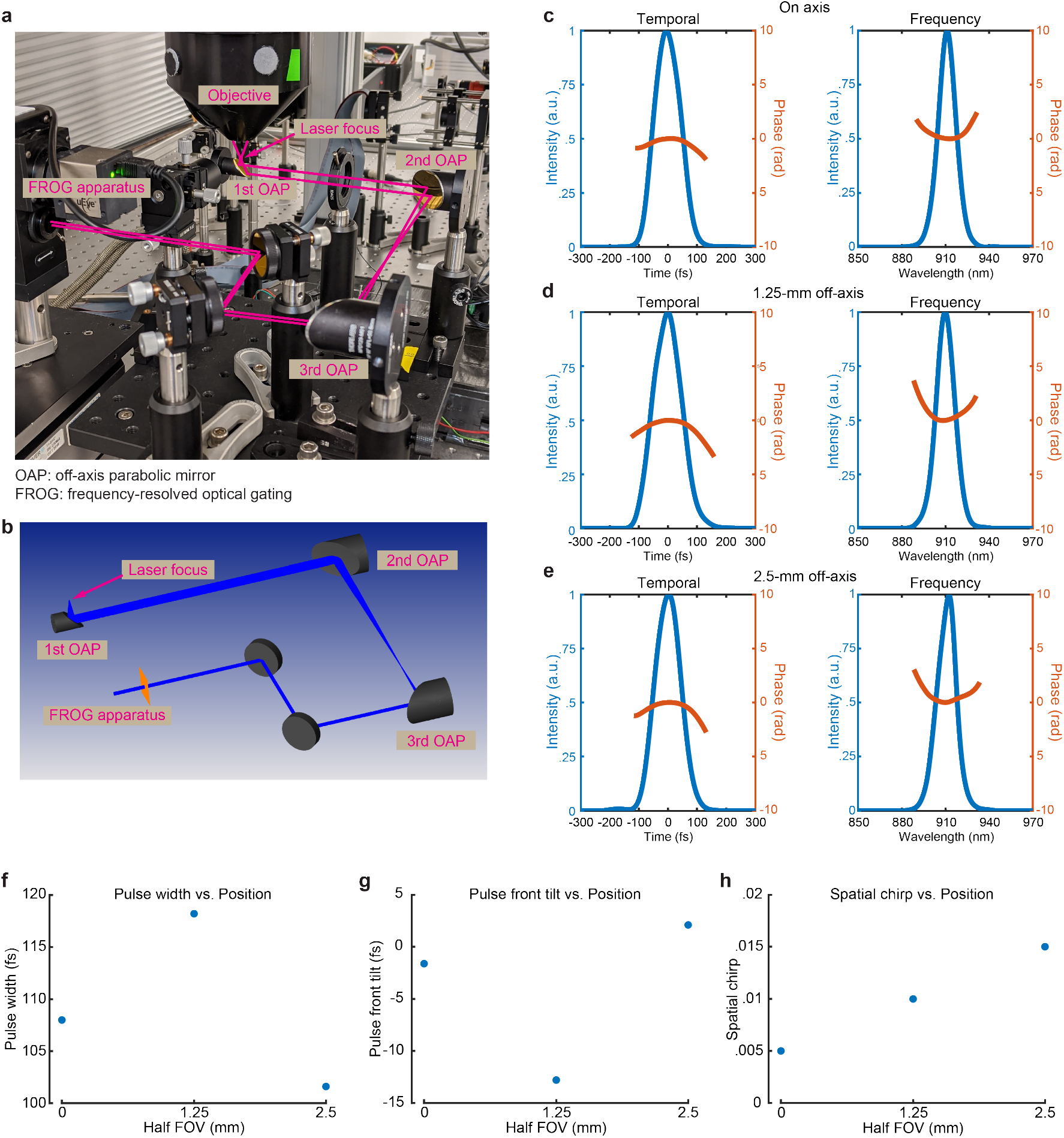
Pulse characterization at the imaging plane. (a) A custom reflective relay redirected light from the focal plane to a pulse metrology device (FROG). (b) Optical model diagram of the beam path. (c-h) The temporal and the spatial measurement of pulses (c) at the center, (d) 1.25-mm off axis, and (e) 2.5-mm off-axis of the FOV. Overall, (f) the pulse width, (g) pulse front tilt, and (h) spatial chirp varied little over the full FOV (mean ± standard error from 5 measurements for each data point).

**Supplementary Figure 10.**
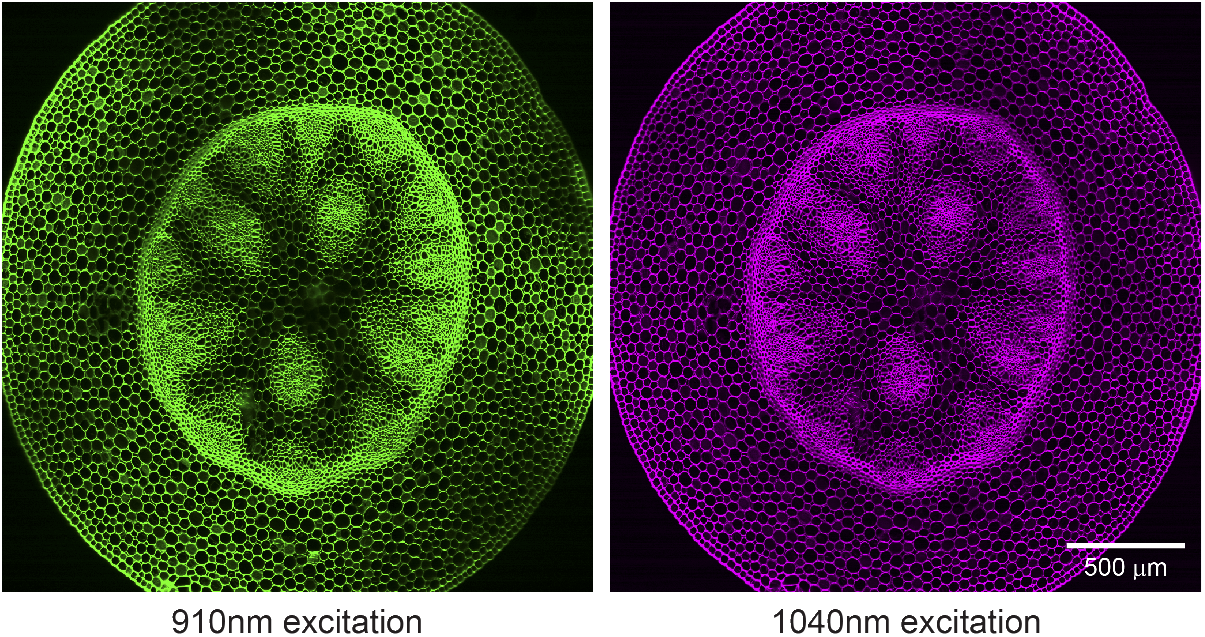
Dual-wavelength large FOV imaging. Two-photon fluorescence images of a fixed convallaria acquired with two excitation wavelengths, 910 nm and 1040 nm, respectively. The image size is 2500 × 2500 µm, acquired with a single raster scan without stitching. The same emission filter, a 700 nm short-pass filter, is used for the two images.

## Video captions

**Supplementary Video 1**

Rapid defocusing using AO. FOV: 375 × 375 µm, acquisition rate: 30 frames/s.

**Supplementary Video 2**

Imaging enhancement using AO. FOV: 659 × 535 µm. acquisition rate: 30 frames/s.

**Supplementary Video 3**

Simultaneous dual region large FOV (1.5 × 5.0 mm each) imaging at high pixel count (1024 × 4096 pixels each). Acquisition rate (3.85 frames/s). Depth: 371 µm.

**Supplementary Video 4**

Simultaneous dual region imaging with flexible imaging configurations. Two regions are 4.36 mm apart. Pathway 1: FOV = 1500 × 1500 µm, acquisition rate = 15 frames/s, pixel count = 1024 × 1024 pixels, depth = 371 µm. Pathway 2: FOV = 256 × 256 µm, acquisition rate = 58 frames/s, pixel count = 256 × 256 pixels, depth = 371 µm.

**Supplementary Video 5**

Simultaneous dual region imaging with part of regions overlapped. Pathway 1: FOV = 250 × 250 µm, acquisition rate = 60 frames/s, pixel count = 256 × 256 pixels, depth = 262 µm. Pathway 2: FOV = 750 × 750 µm, acquisition rate = 15 frames/s, pixel count = 1024 × 1024 pixels, depth = 262 µm.

**Supplementary Video 6**

Quad region imaging with independent parameters and imaging planes. Pathway 1, Region 1 and Region 2: FOV = 1500 × 1500 µm, acquisition rate = 7.6 frames/s, pixel count = 1024 × 1024 pixels, depth = 361 µm. Pathway 2, Region 3 and Region 4: FOV = 250 × 250 µm, acquisition rate = 30 frames/s, pixel count = 256 × 256 pixels, depth = 371 µm.

**Supplementary Video 7**

Random access imaging scan. Each cell body ROI is 21.5 × 21.5 µm, with 16 × 16 pixels. The set of ROIs is acquired 41 times each second, depth = 359 µm.

## Methods

### Optical design and simulations

The entire system including relay, scan, tube, and objective lens systems were modeled in OpticStudio (Zemax, LLC). The subsystems of relay, scan, and tube, and objective lenses were first designed, modeled, and optimized individually. Then, the system was optimized as a whole to further minimize additive aberrations between subsystems. The system was optimized for two wavelengths of 910 ± 10 nm and 1050 ± 10 nm for dual-color imaging in the future. All lenses were custom made by Rocky Mountain Instrument Inc, using tolerances of: radii +/-0.1%, center thickness +/-0.1mm and coated with broad band anti-reflective coating (BBAR), R_avg_ <1.5% at 475nm-1100nm.The relay lenses between the x-resonant scanner and the x-galvo mirror were constructed from the Thorlabs Inc parts (LSM254-1050.ZBB, AC508-250-B-ML). Complete lens data of the Diesel2p system is given in Supplementary Figures 1-5.

### Assembly

The lens sub-assemblies were manufactured, aligned, and assembled in the factory of Rocky Mountain Instrument Co. There are threaded connectors between optical relays connected the two galvo scanners, and between the tube lens and the objective lens. Together with the correction collar on the objective, they are adjustable for the axial separation between subassemblies. Galvo and resonant scanners were mounted on a XY translator (Thorlabs, CXY2), and this was attached to a 60 mm cage cube (Thorlabs, LC6W), which also bridged the subassemblies. As the system is assembled, locations of afocal space at conjugate planes are checked for collimation as designed in the system.

### Animals

All procedures involving living animals were carried out in accordance with the guidelines and regulations of the US Department of Health and Human Services and approved by the Institutional Animal Care and Use Committee at University of California, Santa Barbara. We used GCaMP6s transgenic mice in this study. GCaMP6s transgenic mice were generated by triple crossing of TITL-GCaMP6s mice, Emx1-Cre mice (Jackson Labs stock #005628) and ROSA:LNL:tTA mice (Jackson Labs stock #011008)^21^. TITL-GCaMP6s mice were kindly provided by Allen institute. Mice craniotomy was performed as previously described^22^. Briefly, mice were deeply anesthetized using isoflurane (1.5–2%) augmented with acepromazine (2 mg/kg body weight) during craniotomy surgery. Carpofen (5 mg/kg body weight) was administered prior to surgery, as well as after surgery for 3 consecutive days. A 5-mm diameter cranial window was implanted after removing the scalp overlaying the right visual cortex.

### In vivo two photon imaging

All imaging was performed on the custom Diesel2p system. The instrumentation (see Diesel2p instrumentation below) and image acquisition were controlled by ScanImage from Vidrio Technologies Inc. Animals were awake during calcium imaging. The imaging was performed with <100 mW out of the front of the objective. With typical imaging parameters (512 × 512 at 30 frames/s, 0.5 mm imaging region), no damage was observed from the surface of the dura to the 500 µm depth. Assessment of damage due to laser intensity was based on visual morphological changes to the appearance of the dura mater and/or continuously bright cell bodies.

### Image analysis for neuronal calcium signals

Calcium imaging was processed following our previously established protocol^22^. Ca^2+^ signals were analyzed using custom software in MATLAB (Mathworks). Neurons were segmented and transients were extracted from imaging stacks using custom code adapted from Suite2p modules after motion correction^23^. Calcium transients were reported after neuropil contamination correction that is subtracting the common time series of a spherical surrounding mask of each neuron from the cellular calcium traces. Spike inference was performed using a Markov chain Monte Carlo methods^24^. We computed the Pearson correlation of inferred spike trains between neurons using the MATLAB built-in function *corr*.

### Excitation point spread function measurements and simulations

The measurement and analysis procedure were described in our previous publication in details^11^. To evaluate the excitation point spread function (PSF), sub-micrometer beads were imaged. Sub-micrometer fluorescent beads (0.2 µm, Invitrogen F-8811) were imbedded in a thick (∼1.2 mm) 0.75% agarose gel. 30 µm z-stacks were acquired, each centered at one of three depths (50 µm, 250 µm, 500 µm). The stage was moved axially in 0.5 µm increments (Δ_stage_). At each focal plane 30 frames were acquired and averaged to yield a high signal-to-noise image. Due to the difference between the refractive index of the objective immersion medium (air) and the specimen medium (water), the actual focal position within the specimen was moved an amount Δ_focus_ = 1.38 x Δ_stage_^25^. The factor 1.38 was determined in Zemax and slightly differs from the paraxial approximation of 1.33. These z-stack images were imported into MATLAB for analysis. For the axial PSF, XZ and YZ images were created at the center of a bead, and a line plot was made at an angle maximizing the axial intensity spread, thereby preventing underestimation of the PSF due to tilted focal shifts. For the radial PSF, an XY image was found at the maximum intensity position axially. A line scan in X and Y was made. Gaussian curves were fit to the individual line scans to extract FWHM measurements. The radial PSF values are an average of the X PSF and Y PSF, and the axial PSF is an average of the axial PSF found from the XZ and YZ images. Excitation PSF measurements were performed both on axis and at the edges of the FOV for both imaging pathways. Data reported (Fig. 1i and Supplementary Figure 8b) are the mean ± S.D. of eight beads.

### Diesel2p instrumentation

Our laser source is a Ti:Sapphire pulsed laser, tuned to a central wavelength at 910 nm and a 80 MHz repetition rate (Mai-Tai, Newport). The laser first passes through a built-in pre-chirper unit (DeepSee, Newport), further followed by an external custom-built single prism pre-chirper^15^. The material of the prism is PBH71 (Swamp Optics) with a refractive index of 1.89. The laser power is controlled using a half wave plate (AHWP05M-980, Thorlabs) followed by a polarization beam splitting cube (CCM5-PBS203, Thorlabs). Similar polarization optics were used to split the beam into two paths and control the relative power between the two paths. Prior to splitting, the beam was expanded using a 2× beam expander (GBE02-B, Thorlabs). One beam travels directly to a deformable mirror (DM140A-35-UM01, Boston Micromachines Corporation), and the other beam is first diverted to a delay arm, and subsequently to a separately deformable mirror. The delay arm is designed to impart a 6.25 ns temporal offset to the pulses in one beam (1.875 m additional path length). As the laser pulses are delivered at 12.5 ns intervals (80 MHz), they are evenly spaced in time at 160 MHz after the two beams are recombined. Before the recombination, both pathways pass through their own set of scan engines comprised with a x-resonant scanner (CRS8KHz, Cambridge technology), and a x-galvo scanner (6220H, Cambridge technology), and a y-galvo scanner (6220H, Cambridge technology) in series. These scanners are connected by custom-designed afocal relays. Two beams are recombined with another polarization beam splitter (PC75K095, Rocky Mountain Instrument). A scan lens and tube lens formed a 4× telescope. Together with a short-pass dichroic mirror, they relayed the expanded beams to the back aperture of the custom objective. The entrance scan angles at the objective back aperture were ∼ ± 5 degrees yielding our 5 mm FOV. The generated fluorescence from the imaging plane is directed to a photomultiplier tube (PMT, H10770PA-40 MOD, Hamamatsu) via the assembly of 3 collection lenses (AC508-100-A-ML, AC254-030-A-ML, Thorlabs; 48425, Edmunds Optics). The ultrafast 1040 nm laser used for dual-wavelength imaging was an ALTAIR IR-10 (Spark Lasers).

### Photon counting electronics

Output from the photomultiplier tube was first amplified with a high bandwidth amplifier (C5594-44, Hamamatsu) and then split into two channels (ZFSC-2-2A, Mini-Circuits). One channel was delayed relative to the other by 6.25 ns by using a delay box (DB64, Stanford Research Systems). Each channel was connected to a fast discriminator (TD2000, Fast ComTec GmbH). The ∼80 MHz synchronization output pulses from the laser was delivered to a third fast discriminator (TD2000, Fast ComTec GmbH), which has a continuous potentiometer adjustment to adjust the output pulse width from 1 ns to 30 ns. This output pulse was delivered to the common veto input on the previous two TD2000 discriminators where the PMT outputs were collected. The veto width was adjusted by the potentiometer on third TD2000 discriminator and the relative phase of the veto window was adjusted by delaying the synchronization pulses from the laser module using the DB64 delay box. Outputs from each TD2000 is sent to a channel of the digitizer of the vDAQ card (Vidrio Technologies). Digitized signals were arranged into images with the indicated pixel count in the ScanImage software (Vidrio Technologies). In this manner we could demultiplex the single PMT output into two channels corresponding to the two excitation pathways.

### Pulse characterization at the focal plane

We used the frequency-resolved optical gating (FROG) method measurements to retrieve the pulse conditions at the focal plane using three off-axis parabolic (OAP) mirrors and a FROG system (GRENOUILLE 8-50-334-USB, Swamp Optics). We used reflective OAP mirrors to avoid the post-focus chromatic and spatial dispersion that would be introduced if refractive lenses were used. OAP mirrors help retain more of the original pulse information of the pulses. The focused laser at the focal plane was collimated by the first OAP mirror (MPD00M9-M01, Thorlabs). Then the beam needed to be reduced for measurement. So it was refocused by the second OAP mirror (MPD169-M01, Thorlabs) and re-collimated by a third OAP mirror (MPD129-M01, Thorlabs). At this point, the beam size of the collimated laser was small enough to fit the FROG apparatus’ entrance aperture. The FROG traces were retrieved, and the pulse width, pulse front tilt, and the spatial dispersion were calculated with the built-in retrieval algorithm (QuickFrog, Swamp Optics). The focus was parked at the three FOV locations (on-axis, 1.25-mm off-axis, and 2.5-mm off-axis) for the FROG measurement by deflecting the angle of the X-galvo scanner (0, 5, and 10 degrees), corresponding to the angle of 0, 2.5, and 5.0 degrees off-axis at the entrance pupil of the objective. Rotating the X-galvo scanner (instead of the Y-galvo scanner) maximizes the off-axis traveling pathway for the deflected laser beam in the system, and thus provides an upper-limit to any distortions detected.

## Data availability

The raw calcium imaging data for Figs. 2b and 3 are available at Google Drive: https://drive.google.com/drive/folders/1xjEq9P1M1Km8gzsjodiqRTOxDr0JHzI1?usp=sharing Other data that support the findings of this study are available on request from the corresponding author.

## Code availability

The code used for analysis in this study are available from the corresponding author upon request.

## Author Contributions

CHY assembled, aligned, and optimized the system, collected data, analyzed data, and wrote the manuscript. JNS developed the concept, designed the optics, oversaw manufacturing, and collected pilot data. YY performed mouse surgeries, consulted on imaging, and analyzed data. RH optimized the system and collected pilot data. SLS supervised the project.

## Acknowledgements

We thank Ikuko Smith for scientific mentorship and lab leadership. We thank Swamp Optics for the custom prism. This work was supported by the McKnight Foundation, Human Frontier Science Program (RGP0027/2016), NSF (NeuroNex #1934288 and BRAIN EAGER #1450824) and the NIH (NINDS R01NS091335 and NEI R01EY024294).

## Notes

### Competing Interest Statement

The authors have declared no competing interest.

## Reference

1 Denk, W., Strickler, J. H. & Webb, W. W. Two-photon laser scanning fluorescence microscopy. Science 248, 73, doi: 10.1126/science.2321027 (1990).

2 Chen, J. L., Andermann, M. L., Keck, T., Xu, N. L. & Ziv, Y. Imaging Neuronal Populations in Behaving Rodents: Paradigms for Studying Neural Circuits Underlying Behavior in the Mammalian Cortex. Journal of Neuroscience 33, 17631–17640, doi: 10.1523/Jneurosci.3255-13.2013 (2013).

3 Sofroniew, N. J., Flickinger, D., King, J. & Svoboda, K. A large field of view two-photon mesoscope with subcellular resolution for in vivo imaging. Elife 5, doi: 10.7554/eLife.14472 (2016).

4 Ota, K. et al. Fast scanning high optical invariant two-photon microscopy for monitoring a large neural network activity with cellular resolution. bioRxiv, 2020.2007.2014.201699, doi: 10.1101/2020.07.14.201699 (2020).

5 Bumstead, J. R. et al. Designing a large field-of-view two-photon microscope using optical invariant analysis. Neurophotonics 5, 025001, doi: 10.1117/1.NPh.5.2.025001 (2018).

6 Ji, N., Freeman, J. & Smith, S. L. Technologies for imaging neural activity in large volumes. Nat Neurosci 19, 1154–1164, doi: 10.1038/nn.4358 (2016).

7 Amir, W. et al. Simultaneous imaging of multiple focal planes using a two-photon scanning microscope. Opt. Lett. 32, 1731–1733, doi:Doi 10.1364/Ol.32.001731 (2007).

8 Rumyantsev, O. I. et al. Fundamental bounds on the fidelity of sensory cortical coding. Nature 580, 100–105, doi: 10.1038/s41586-020-2130-2 (2020).

9 Cheng, A., Goncalves, J. T., Golshani, P., Arisaka, K. & Portera-Cailliau, C. Simultaneous two-photon calcium imaging at different depths with spatiotemporal multiplexing. Nature Methods 8, 139–U158, doi: 10.1038/Nmeth.1552 (2011).

10 Zhang, T. et al. Kilohertz two-photon brain imaging in awake mice. Nature Methods 16, 1119-+, doi: 10.1038/s41592-019-0597-2 (2019).

11 Stirman, J. N., Smith, I. T., Kudenov, M. W. & Smith, S. L. Wide field-of-view, multi-region, two-photon imaging of neuronal activity in the mammalian brain. Nat Biotechnol 34, 857–862, doi: 10.1038/nbt.3594 (2016).

12 Chen, J. L., Voigt, F. F., Javadzadeh, M., Krueppel, R. & Helmchen, F. Long-range population dynamics of anatomically defined neocortical networks. Elife 5, doi: 10.7554/eLife.14679 (2016).

13 Tsyboulski, D. et al. Remote focusing system for simultaneous dual-plane mesoscopic multiphoton imaging. bioRxiv, 503052, doi: 10.1101/503052 (2018).

14 Yang, W. et al. Simultaneous Multi-plane Imaging of Neural Circuits. Neuron 89, 269–284, doi: 10.1016/j.neuron.2015.12.012 (2016).

15 Akturk, S., Gu, X., Kimmel, M. & Trebino, R. Extremely simple single-prism ultrashort-pulse compressor. Opt. Express 14, 10101–10108, doi:Doi 10.1364/Oe.14.010101 (2006).

16 Trebino, R. et al. Measuring ultrashort laser pulses in the time-frequency domain using frequency-resolved optical gating. Review of Scientific Instruments 68, 3277–3295, doi: 10.1063/1.1148286 (1997).

17 Chen, T.-W. et al. Ultrasensitive fluorescent proteins for imaging neuronal activity. Nature 499, 295–300, doi: 10.1038/nature12354 (2013).

18 Lu, R. W. et al. Rapid mesoscale volumetric imaging of neural activity with synaptic resolution. Nature Methods 17, 291-+, doi: 10.1038/s41592-020-0760-9 (2020).

19 Beaulieu, D. R., Davison, I. G., Kılıç, K., Bifano, T. G. & Mertz, J. Simultaneous multiplane imaging with reverberation two-photon microscopy. Nature Methods 17, 283–286, doi: 10.1038/s41592-019-0728-9 (2020).

20 Yang, W. & Yuste, R. Holographic imaging and photostimulation of neural activity. Curr Opin Neurobiol 50, 211–221, doi: 10.1016/j.conb.2018.03.006 (2018).

21 Madisen, L. et al. Transgenic Mice for Intersectional Targeting of Neural Sensors and Effectors with High Specificity and Performance. Neuron 85, 942–958, doi: https://doi.org/10.1016/j.neuron.2015.02.022 (2015).

22 Yu, Y., Stirman, J. N., Dorsett, C. R. & Smith, S. L. Mesoscale correlation structure with single cell resolution during visual coding. bioRxiv, 469114, doi: 10.1101/469114 (2018).

23 Pachitariu, M. et al. Suite2p: beyond 10,000 neurons with standard two-photon microscopy. bioRxiv, 061507, doi: 10.1101/061507 (2017).

24 Pnevmatikakis, E. A., Merel, J., Pakman, A. & Paninski, L. in 2013 Asilomar Conference on Signals, Systems and Computers. 349–353.

25 Visser, T. D. & Oud, J. L. Volume measurements in three-dimensional microscopy. Scanning 16, 198–200, doi: 10.1002/sca.4950160403 (1994).

